# Lymphangiogenesis is Critical for Healing and Survival in a Murine Model of Laryngotracheal Injury

**DOI:** 10.64898/2026.08.19.745806

**Authors:** Peter Gulleman, Yueli Zhang, Finnegan Clark, Maxim Litvak, Alexander Clinton, Alexander Hillel, Gail Deutsch, Tzushan Sharon Yang, Alexander Gelbard, Jennifer M Sucre, Jason S Park

## Abstract

**Objective:** Lymphatic dysfunction has been implicated in exacerbating fibrosis in numerous diseases, yet the role of the lymphatic system in laryngotracheal injury has not previously been explored. This study aims to evaluate lymphatic vascular remodeling in a murine model of laryngotracheal stenosis (LTS) and determine how pharmacologic blockade of lymphangiogenesis impacts airway healing after mucosal injury.

**Methods:** LTS was induced in C57BL6 mice using an established chemomechanical injury model. Lymphatic density was quantified using LYVE-1 immunohistochemistry. Mice were treated with the VEGFR-3-selective tyrosine kinase inhibitor SAR131675 to block lymphangiogenesis after injury. Outcomes assessed included survival, histopathology, immunohistochemistry, and Evans blue dye vascular leakage.

**Results:** Laryngotracheal injury induced a substaantial increase in subepithelial lymphatic vessel density concomitant with fibrotic remodeling. Pharmacologic inhibition of VEGFR-3 signaling with SAR131675 abrogated this lymphangiogenic response and resulted in markedly increased mortality, impaired epithelial repair with obstructive sloughing, increased edema, and persistent histopathologic evidence of tissue injury. A qualitative increase in pathologic fibrocellular remodeling was also observed, though with no measurable difference in lamina propria thickness.

**Conclusion:** These findings establish lymphatic remodeling as an essential component of successful airway repair following mucosal injury. Lymphatic dysfunction is a common feature of known risk factors for LTS including diabetes, obesity, and prematurity, and can be exacerbated by positive pressure ventilation. Disruption of the lymphangiogenic response to airway injury may lead to stasis of pro-inflammatory factors that result in chronic inflammation, maladaptive remodeling, and pathologic tissue changes. The lymphatic vasculature is a viable target for future mechanistic study and potential therapeutic intervention following airway injury.

## Introduction

Laryngotracheal stenosis (LTS) is a life-threatening fibroproliferative upper airway narrowing, most commonly arising from iatrogenic mucosal injury and characterized by reactive inflammation, profibrotic cytokine production, and fibroblast proliferation^1^. Inflammatory edema and ultimately excessive collagen deposition from activated fibroblasts results in obstructive airway scar formation^2–6^. Iatrogenic LTS most commonly arises as a sequela of prolonged endotracheal intubation or tracheostomy. Limited understanding of the molecular mechanisms underlying LTS has impeded attempts to develop effective medical therapies for the condition. Current treatments are therefore primarily surgical, underscoring the critical need for novel therapeutic strategies that target the underlying mechanisms of airway fibrosis^7^.

The lymphatic system is composed of lymphoid organs (including lymph nodes) and a network of endothelium-lined vessels that course throughout the body^8^. A functional lymphatic system is crucial for the balance and transport of interstitial fluid, antigens, and macromolecules as well as for regulation of immune cell trafficking. Efficient lymphatic drainage is essential for the resolution of inflammatory processes through removal of edema fluid and facilitating the egress of immune cells and inflammatory mediators to regional lymph nodes^9^. Lymphatic dysfunction, particularly dysregulated lymphangiogenesis, has been implicated in exacerbating fibrosis in various diseases including pulmonary fibrosis, organ transplant rejection, cardiac fibrosis, renal fibrosis, liver fibrosis, and skin fibrosis^10–14^.

Although well-described in other disease processes, the role of the lymphatic system in laryngotracheal injury has not been explored. To tackle this knowledge gap, we used an established murine model and targeted the vascular endothelial growth factor receptor 3 (VEGFR-3) signaling pathway^5,15–17^. VEGFR-3, expressed on all mature lymphatic endothelial cells (LEC) and a primary driver of lymphatic growth and function^18,19^, can be selectively blocked by the orally available VEGFR- 3 tyrosine kinase inhibitor SAR131675^20^. We found that inhibiting lymphangiogenesis impaired tracheal repair after injury, suggesting that peri-injury lymphatic expansion is compensatory rather than pathologic, underpinning a new paradigm for future treatment.

## Results

### Murine Laryngotracheal Injury Induces Lymphangiogenesis

To investigate how laryngotracheal injury impacts the tracheal lymphatic vasculature, we utilized an established murine model of laryngotracheal stenosis initiated by chemomechanical trauma using a bleomycin-dipped wire brush (**Figure 1A**). Control mice underwent sham surgery with cervical skin incision only.

**Figure 1.**
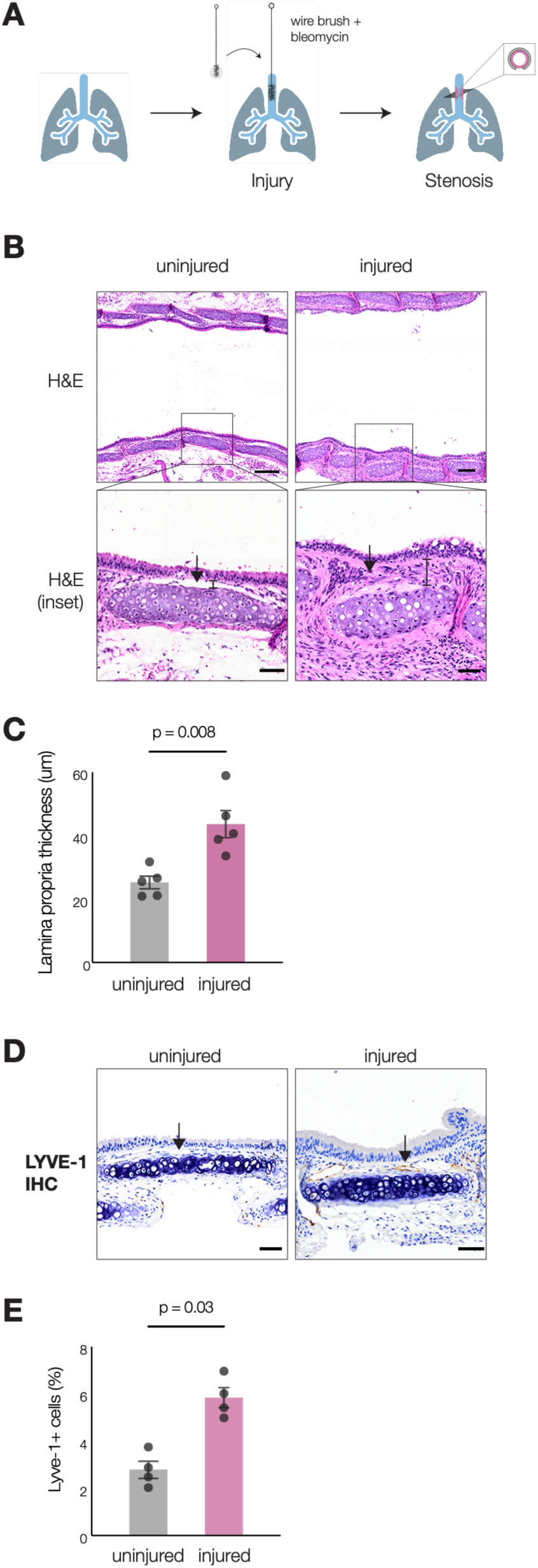
Murine laryngotracheal injury causes a significant increase in subepithelial lymphatic vessels concomitant with fibrosis development. **1A** - Schematic of the chemomechanical laryngotracheal injury model. After the laryngotracheal complex is exposed through a cervical incision for visualization and counter-traction, the tracheal mucosa is abraded with a bleomycin-dipped wire brush inserted through a transorally inserted intratracheal catheter. **1B** - Representative H&E-stained tracheal tissue sections from uninjured and injured mice 2 weeks after injury, demonstrating characteristic features of post-injury fibrotic remodeling including epithelial flattening, decreased ciliated epithelium, squamous metaplasia, and thickening of the lamina propria (arrows and dimension line). Scale bar for top images = 200 μm. Scale bar for zoomed insets = 50 μm. **1C** - Quantification of lamina propria thickness shows significant increase following laryngotracheal injury, bar graph. p = 0.008 by Mann-Whitney U test. Error bars represent mean ± SEM. **1D** - Representative immunohistochemical staining for LYVE-1 showing increased lymphatic vessel density in the lamina propria of injured tracheas compared to uninjured controls (arrows). Brown DAB chromogen, anti-LYVE-1. Scale bar = 50 μm. **1E** - Quantification of LYVE-1+ cells as a percentage of subepithelial area demonstrates a significant increase following injury. *, p = 0.03 by Mann-Whitney U test. Error bars represent mean ± SEM.

We first confirmed the expected histologic features of fibrosis by comparing injured and uninjured airway tissue 2 weeks after laryngotracheal injury. Histological analysis of injured tracheal tissue revealed lamina propria (LP) thickening by matrix deposition, epithelial flattening, a decrease in ciliated epithelium, and squamous metaplasia on hematoxylin and eosin (H&E) staining (**Figure 1B**). Stenosis was quantified by measuring LP thickness, with injured airways showing an average of 43.7 μm (95CI: 31.7-55.7 μm) and uninjured airways an average of 25.2 μm (95CI: 19.6-30.8 μm) (p=0.008) (**Figure 1C**). These results were consistent with previously published descriptions of post- injury fibrotic remodeling in this injury model. Next, we investigated whether lymphatic vessel density is altered after laryngotracheal injury by performing immunohistochemical staining for the lymphatic endothelial cell marker LYVE-1 (lymphatic vessel endothelial hyaluronan receptor 1) (**Figure 1D**).

This revealed a significant increase in LECs within the injured trachea as compared to uninjured controls, from 2.8% (95CI: 1.6%-4.0%) to 5.8% (95CI: 4.5%-7.2%) of submucosal area (p = 0.03), indicating a robust lymphangiogenic response to injury (**Figure 1E**).

### VEGFR-3 Inhibition Worsens Survival and Abrogates the Lymphangiogenic Response to Laryngotracheal Injury

Next we investigated how VEGFR3 inhibition impacts the tracheal tissue response to laryngotracheal injury. Mice were treated daily with SAR131675, a highly selective VEGFR-3 tyrosine kinase inhibitor, starting postoperative day 1 after laryngotracheal injury (**Figure 2A**). Treatment with SAR131675 (as compared to vehicle control) significantly reduced survival in mice that underwent laryngotracheal injury, with death occurring in 54% of treated mice by 7 days after injury (p = 0.004) (**Figure 2B**). Uninjured mice that underwent treatment with SAR131675 without laryngotracheal injury did not show a difference in survival as compared to vehicle control (p > 0.99) (**Figure 2B**).

**Figure 2.**
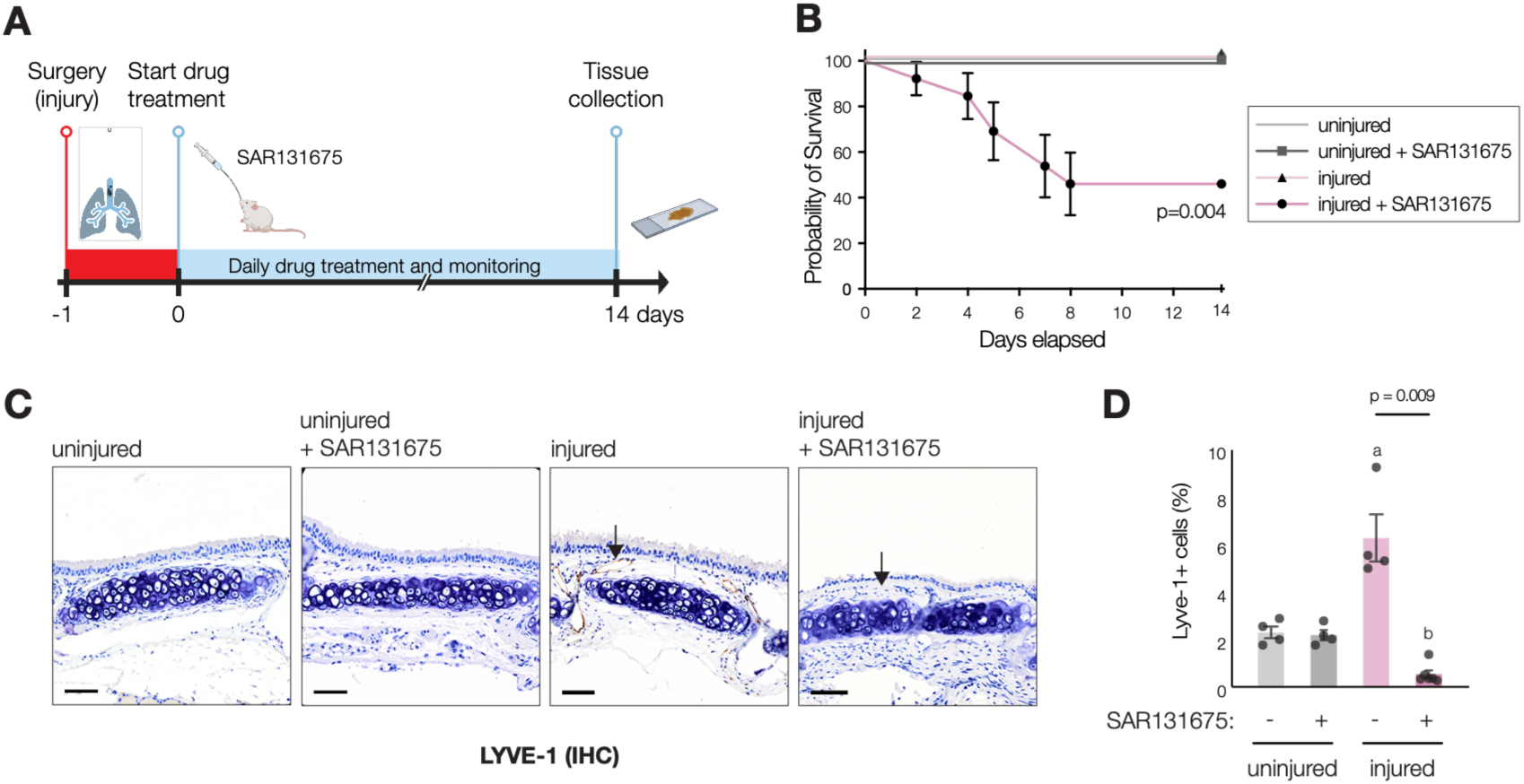
Blockade of lymphangiogenesis through systemic administration of the selective VEGFR-3 tyrosine kinase inhibitor SAR131675 after laryngotracheal injury abrogates the lymphangiogenic response and decreases survival. **2A** - Experimental timeline for oral SAR131675 treatment following laryngotracheal injury. Drug (100 mg/kg) or vehicle control (corn oil) was administered by daily oral gavage starting postoperative day 1. **2B** - Mice treated with SAR131675 after laryngotracheal injury exhibit significantly reduced survival compared to vehicle-treated controls, with mortality occurring predominantly within the first 7 days post-injury. Kaplan-Meier survival curve. p = 0.004 by log-rank (Mantel-Cox) test. **2C** - Representative LYVE-1 immunohistochemical staining of tracheal tissue collected 14 days after injury. Tracheas from injured vehicle-treated mice show robust lymphatic vessel expansion in regions of scar, whereas tracheas from SAR131675-treated mice show essentially complete absence of lymphatic vessels (arrows). Uninjured mouse tracheas show relatively few lymphatics in both SAR131675 drug-treated and untreated conditions. Scale bar = 50μm. **2D** - Quantification of LYVE-1+ cells as a percentage of subepithelial area in panel 2C, bar graph. SAR131675 treatment abrogated injury-induced lymphangiogenesis (p = 0.009), with lymphatic density falling below that observed in uninjured controls. p = 0.009 by Mann-Whitney U test. SAR131675 treatment alone did not alter lymphatic density in uninjured tissues (ns, p > 0.99). a, LYVE-1+ cell density was also higher in injured vehicle-treated mice compared to uninjured vehicle-treated mice (p = 0.03) and uninjured SAR131675-treated mice (p = 0.03). b, LYVE-1+ cell density was also lower in injured SAR131675-treated mice compared to uninjured vehicle-treated mice (p = 0.01) and uninjured SAR131675-treated mice (p = 0.01).

We then performed immunohistochemical staining for LYVE-1 to investigate the impact of VEGFR-3 inhibition on the previously observed expansion in tissue lymphatic vessel density after injury. Tissue specimens were collected 14 days after injury along with vehicle-treated controls (**Figure 2C**). In stark contrast to the increase in lymphatic density normally observed after injury (**Figure 1D**), treatment with SAR131675 essentially eliminated lymphatic vessels in regions of injury: 6.3% (95CI: 3.1%-9.5%) vs 0.54% (95CI: 0.1%-1.0%) (p = 0.009) (**Figure 2D**). SAR131675 treatment did not alter lymphatic density in uninjured tissues as compared to treatment with vehicle control: 2.2% (95CI: 1.5%-2.9%) vs 2.3% (95CI: 1.5%-3.1%) (p = 0.99) (**Figure 2D**). Notably, lymphatic density in injured SAR131675- treated mice was also statistically significantly lower than the baseline lymphatic density found in tissue from uninjured mice (p = 0.01).

### Blockade of Lymphangiogenesis Increases Vascular Leakage and Tissue Edema

We investigated the physiologic mechanisms by which VEGFR-3 inhibition may be leading to death after injury. To determine whether disruption of lymphangiogenesis may be impairing clearance of interstitial fluid and inflammatory mediators, we used an Evans Blue dye extravasation assay to quantify vascular leakage and tracheal tissue edema. Mice were sacrificed 5 days after injury to assess mice prior to the time window death was observed in the prior experiment.

Tracheas from injured mice treated with SAR131675 showed an increase in normalized tissue Evans Blue accumulation, 7.0 (95CI: 4.5-9.6) units/mg with SAR131675-treatment vs 4.6 (95CI: 3.2-6.0) units/mg with vehicle control (p=0.03) (**Figures 3A-3B**), consistent with impaired lymphatic-mediated fluid clearance and accumulation of inflammation-associated tissue edema. There was no difference in Evans Blue accumulation in uninjured mice treated with SAR131675 as compared to vehicle control treatment: 4.6 (95CI: 4.1-5.1) units/mg with SAR131675 vs 4.4 (95CI: 3.4-5.5) units/mg with vehicle control (**Figure 3B**).

**Figure 3.**
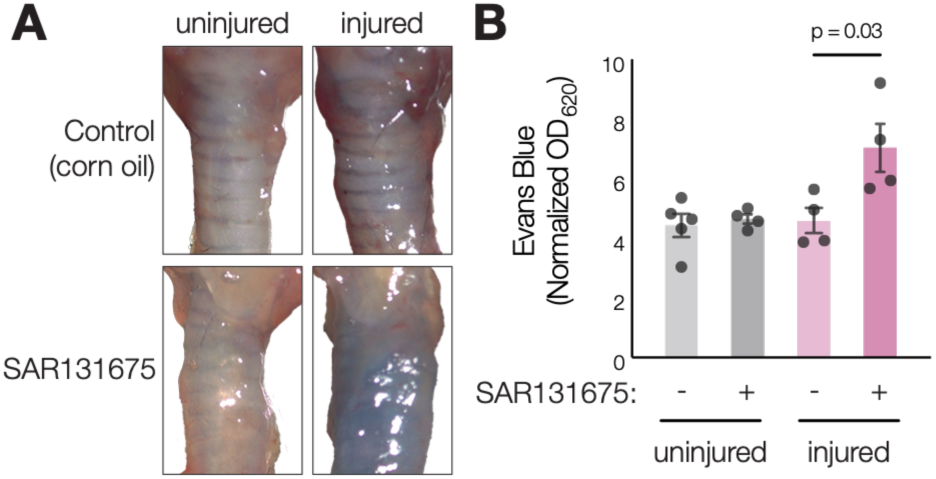
Inhibition of lymphangiogenesis by SAR131675 treatment after laryngotracheal injury increases vascular leakage and tissue edema. **3A** - Representative photomicrographs of tracheal tissue following retro-orbital Evans Blue dye perfusion demonstrate increased dye extravasation in tracheas from SAR131675-treated mice compared to vehicle-treated controls. Each image is approximately 6mm wide by 9mm tall and includes the laryngeal complex and about 7 tracheal rings. **3B -** Quantification of Evans Blue dye content from tracheal tissue using formamide extraction and spectrophotometry. OD620 readings normalized to tissue wet weight demonstrate elevated dye accumulation in tracheas from SAR131675-treated mice following laryngotracheal injury (p = 0.03 by Mann-Whitney U test). No significant difference was observed between treatment groups in uninjured mice.

Inhibition of Lymphangiogenesis Exacerbates Histopathologic Evidence of Airway Injury We hypothesized that accumulation of inflammatory mediators following lymphatic impairment would disrupt airway repair. First, comparison of LP thickness between SAR131675- and vehicle-treated injured mice revealed no statistically significant difference: 42.4 μm (95CI: 32.8-52.0 μm) for SAR131675 versus 44.0 μm (95CI: 29.7-58.4 μm) for vehicle control (p=0.90) (**Figure 4A**). Additionally, Picro Sirius Red staining – which detects collagen – also revealed no statistically significant difference in stain-positive area: 84% (95CI: 0.77-0.90) for SAR131675 versus 83% (95CI: 0.63-1.02) (p>0.99) (**Figure 4B**).

**Figure 4.**
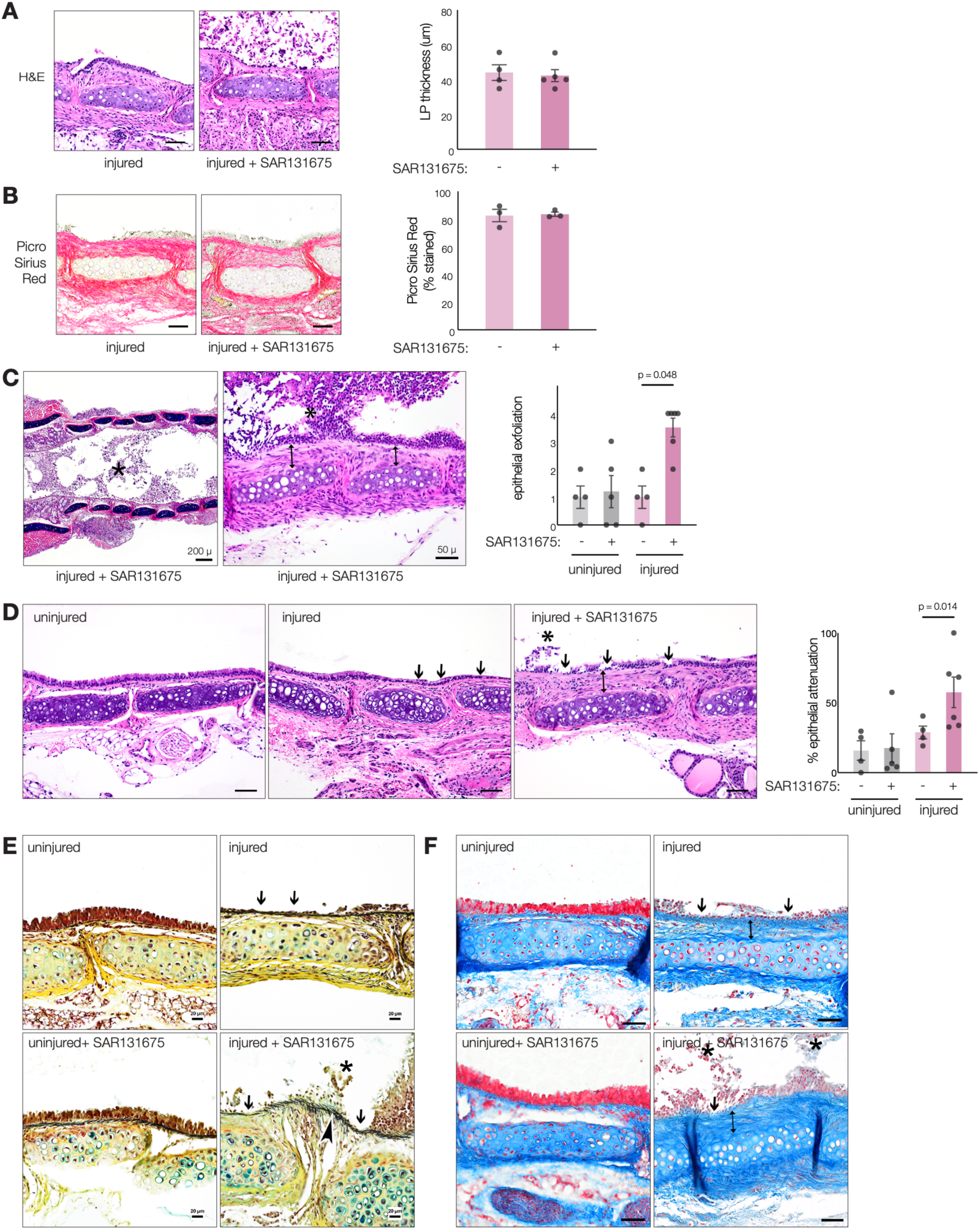
Inhibition of lymphangiogenesis by SAR131675 treatment after laryngotracheal injury worsens histopathologic evidence of airway injury. **4A** - Representative H&E-stained tracheal sections from mice treated with vehicle versus SAR131675 after airway injury. Scale bar = 50 μm. SAR131675-treated mice show worsened epithelial barrier architecture and cellular sloughing into the airway lumen, but no statistically significant difference in lamina propria thickness: 42.4 μm (95CI: 32.8-52.0 μm) for SAR131675 versus 44.0 μm (95CI: 29.7-58.4 μm) for vehicle control p=0.90). **4B** - Representative Picro Sirius Red-stained tracheal sections from mice treated with vehicle versus SAR131675 after airway injury. Scale bar = 50 μm. Bar graph shows quantification of stain-positive area with no statistically significant difference between groups: 84% (95CI: 0.77-0.90) for SAR131675 versus 83% (95CI: 0.63-1.02) (p>0.99). **4C** - Representative H&E-stained tracheal sections from a mouse treated with SAR131675 after airway injury showing epithelial exfoliation and sloughing with accumulation of obstructive debris within the airway lumen (*) as well as subepithelial fibrosis (dimension lines). Bar graph shows pathologist quantification of epithelial exfoliation (0–4 score) demonstrating a statistically significant increase over vehicle-treated controls. Scale bars = 200 μm (low magnification), 50 μm (high magnification). **4D** - Representative H&E-stained tracheal sections demonstrating areas of attenuated or denuded epithelium (arrows), as well as subepithelial fibrosis (dimension line). Bar graph shows pathologist quantification of epithelial attenuation (0-100% score) showing significantly increased epithelial attenuation in comparison with vehicle-treated mice. Scale bars = 50 μm. **4E** - Representative modified Movat’s pentachrome staining demonstrates increased dense subepithelial fibrocellular remodeling and more robust disorganized elastic fiber deposition (arrowhead) in tracheas from SAR131675-treated mice after injury as compared to vehicle-treated and uninjured controls, in addition to epithelial attenuation (small arrows) and sloughing (*). Scale bar = 20 μm. **4F** - Representative Martius Scarlet Blue trichrome staining also demonstrates epithelial attenuation (small arrows) and subepithelial fibrosis (dimension line). Fibrosis extends beyond tracheal cartilage and sloughing and accumulation of epithelial cells in the airway lumen (*) is present in SAR131675- treated mice after injury. Scale bar = 50 μm.

We next performed a detailed histopathologic analysis. Longitudinal tracheal sections were examined by two board-certified pathologists (T.S.Y., veterinary; G.D., clinical), comparing tissue sections from mice treated with SAR131675 or with vehicle control after laryngotracheal injury. On H&E stains, tracheal tissues from mice treated with SAR131675 showed substantial epithelial exfoliation with obstructive sloughing of cellular debris into the airway lumen: pathologist score of 3.5 (95CI: 2.6-4.4) for SAR131675 versus 1.0 (95CI: −0.3-2.3) for vehicle (p=0.014) (**Figure 4C**). In addition, SAR131675-treated injured mice showed significantly worsened epithelial attenuation in comparison with vehicle-treated mice: 57.6% (95CI: 29.4%-85.7%) for SAR131675 versus 28.9% (95CI: 14.5%- 43.2%) for vehicle (p=0.048) (**Figure 4D**). There were no statistically significant differences in either measure for uninjured mice treated with SAR131675 versus vehicle.

Finally, Martius Scarlet Blue trichrome and modified Movat’s pentachrome staining was used for additional evaluation of the extracellular matrix architecture by a pathologist (G.D.). In comparison with vehicle controls, tracheas from injured SAR131675-treated mice showed a qualitatively increased severity of subepithelial fibrocellular remodeling as well as persistent inflammation and more robust, disorganized elastic fiber deposition (**Figure 4E-4F**).

### Methodology

#### Murine laryngotracheal injury model

We utilized an established, well-validated murine model of LTS involving combined chemomechanical injury to the laryngotracheal complex^17^. Both male and female adult C57BL/6 mice aged at least 9 weeks (female) or 7 weeks old (male) were used. Briefly, ketamine (100 mg/kg) and xylazine (5 mg/kg) anesthetized mice underwent cervical incision and exposure of the laryngotracheal complex for visualization and counter-traction, followed by transoral intubation with a catheter through which a bleomycin-soaked wire brush abraded the tracheal mucosa. Mice were monitored for humane endpoints until sacrifice at 5 days (Evans Blue experiments) or 2 weeks. All procedures were approved by the Vanderbilt Institutional Animal Care & Use Committee and adhered strictly to institutional and U.S. national guidelines for animal welfare and research ethics.

#### Pharmacologic inhibition of lymphangiogenesis

SAR131675, a selective VEGFR3 tyrosine kinase inhibitor, was purchased from TargetMol Chemicals Incorporated (Boston, MA) (item T6012). The drug was administered orally by gavage to mice daily starting one day after laryngotracheal injury (100 mg/kg/day, corn oil diluent, total administered volume 200 μL).

#### Evans blue dye extravasation assay

Tissue vascular permeability was assessed by Evans blue dye extravasation^21^. Evans Blue (200 μL of 0.5% in PBS) was administered by retro-orbital injection 30 minutes prior to sacrifice. Quantification was performed by dye extraction from tracheal tissue using formamide followed by spectrophotometric analysis at 620nm. OD620 readings were normalized to tissue wet weight.

#### Histopathology

Tracheas were fixed in 10% neutral buffered formalin, processed, paraffin-embedded, then sectioned using a microtome. Tissues were embedded in longitudinal orientation to visualize the entire length of the trachea including the zone of injury. Slides were stained with hematoxylin and eosin (H&E), anti- LYVE-1 (Angiobio 11034) for immunohistochemistry, Picro Sirius Red (Abcam ab245887), Martius Scarlet Blue, and Movat Pentachrome (Abcam ab245884). Blinded measurements of lamina propria (LP) thickness were performed from tracheal rings 2-6 using open-source ImageJ and QuPath software. Quantification of IHC and colorimetric staining was performed using QuPath software.

Sections were evaluated by two board-certified pathologists (G.D., clinical; T.S.Y., veterinary). Sections were scored blindly (T.S.Y.) for epithelial integrity, epithelial exfoliation/sloughing, LP thickness, inflammation, and edema. Epithelial injury was quantified as described above. Edema and inflammation were scored on severity (0 = normal, 1= minimal, 2 = mild, 3 = moderate, 4 = severe). Epithelial exfoliation score (0-4) was assigned by number of sloughed cells within the tracheal lumen (<5, 5–10, 10–30, 30–50, and >50 cells, respectively).

### Statistical analysis

Statistical testing was performed in GraphPad Prism. Mann-Whitney U test (unpaired, non-parametric) was used for two-group comparisons and the log-rank (Mantel-Cox) test for survival analysis. Significance was defined as p < 0.05.

## Discussion

### Summary of Key Findings

We demonstrate that lymphangiogenesis is critical in the early tissue response to laryngotracheal injury. Injury induced a robust expansion of subepithelial lymphatic vessels and VEGFR-3 inhibition with SAR131675 abrogated this response, resulting in markedly increased mortality, impaired epithelial repair with obstructive sloughing, increased tissue edema, and persistent histopathologic evidence of tissue injury. Interestingly, inhibition of lymphangiogenesis qualitatively increased the histopathologic severity of fibrocellular remodeling though without a detectable increase in LP thickness or Picro Sirius Red stain-positive area at the two-week timepoint. Thus the principal consequence of impaired lymphatic remodeling during early wound healing, likely cause of mortality in this study, is defective resolution of acute injury rather than accelerated fibrosis.

Our results support a mechanistic framework in which the lymphatic system remodels acutely to clear inflammatory mediators and immune cells, enabling subsequent repair^11,22,23^. In this model, disrupting lymphangiogenesis causes stasis of pro-inflammatory factors leading to chronic inflammation and pathologic remodeling (**Figure 5**). Lymphatic expansion facilitates clearance of inflammatory edema, cytokines, danger-associated molecular patterns, and activated immune cells from injured tissue, thereby permitting tissue repair and restoration of epithelial integrity. VEGFR-3 inhibition aborts this adaptive response, resulting in the failure of inflammatory resolution. Consistent with contemporary concepts in resolution biology in which successful wound healing depends upon active termination of inflammation rather than passive decay^57–61^, these findings suggest that lymphatic function contributes to the restoration of tissue homeostasis after airway injury.

**Figure 5.**
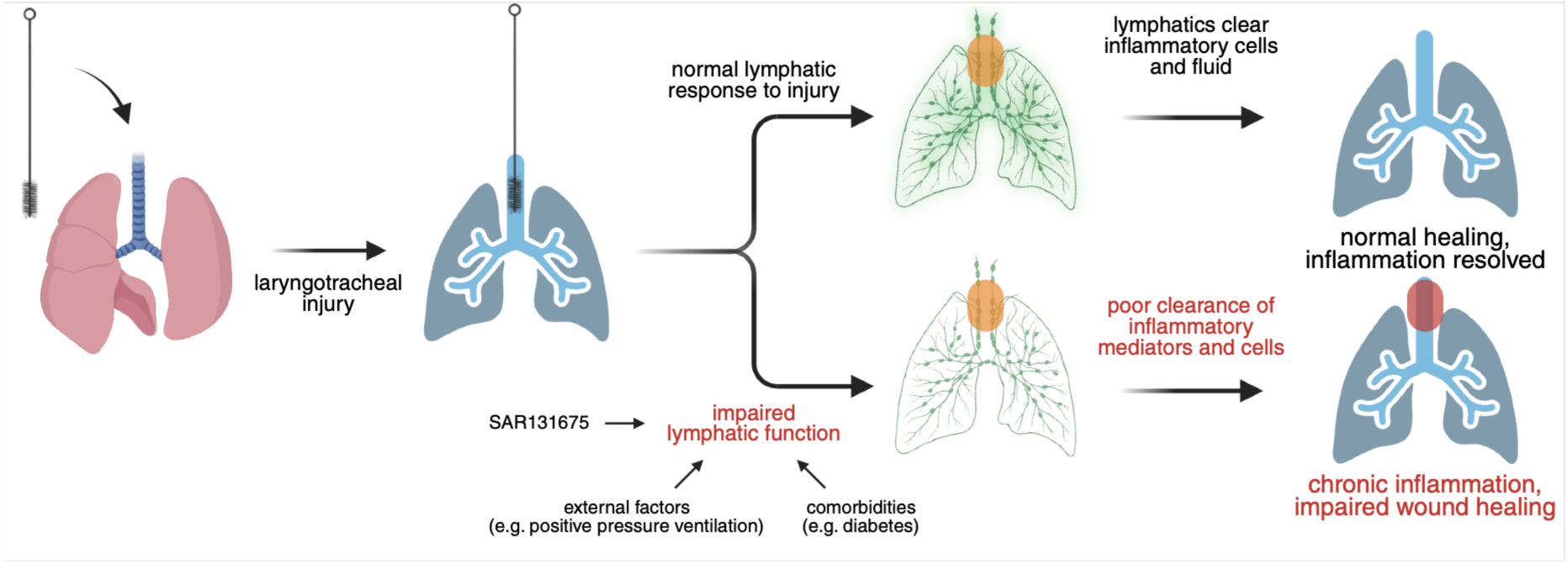
Proposed mechanistic model. Following laryngotracheal mucosal injury, injury-induced lymphangiogenesis is necessary to resolve inflammatory edema and clear immune cells and inflammatory mediators and set the stage for normal wound healing. Disruption of lymphangiogenesis leads to a pathologic buildup of inflammatory mediators and interstitial fluid culminating in persistent inflammation, impaired wound healing with persistent epithelial barrier dysregulation, and increased mortality. Created in BioRender. Park, J. (2026) https://BioRender.com/xirqlpg

### Strengths and Limitations

This study’s principal strength is the integration of multiple approaches including histopathology, quantitative lymphatic vessel analysis, physiologic assessment of vascular permeability, and survival to evaluate a novel concept - the role of lymphangiogenesis during airway repair. These data demonstrate concordant effects that support a meaningful role for lymphatic remodeling in the airway injury response. The chemomechanical injury model closely recapitulates key pathological features of human laryngotracheal stenosis, increasing translational relevance^24^.

Several limitations should also be acknowledged. First, pharmacologic inhibition of VEGFR3 signaling disrupts only one aspect of lymphatic biology – VEGFR3-dependent lymphangiogenesis. It does not impair lymphatic function directly. Also, while SAR131675 is highly selective for VEGFR3 (which is predominantly expressed on LECs), VEGFR3 signaling inhibition could influence other biological processes, and SAR131675 could have off-target effects not previously described. Next, LYVE-1 immunostaining quantifies lymphatic vessel abundance but does not directly measure lymphatic function. Future studies employing functional lymphatic drainage assays will be important to evaluate the physiologic function of newly formed lymphatic vessels observed after laryngotracheal injury.

Finally, although increased Evans blue accumulation is consistent with impaired clearance of tissue edema, the downstream inflammatory mediators and immune-cell populations responsible for defective tissue repair remain an important future direction for this research.

We note that LP thickness values in the present work show small numerical differences compared to previously published work by other labs with this injury model. Reported uninjured LP thickness ranges from 14.5μm to 40.8 μm (present study: 25.2μm). Injured LP thickness ranges from 45.3μm to 95.9μm (present study: 43.7μm). Uniquely in our work we section the trachea longitudinally to visualize and quantitate stenosis from tracheal rings 2-6 to obtain an unbiased average LP thickness measurement across the region. This expectedly results in a lower average LP thickness than measuring thickness in select transverse-cut stenotic tracheal ring sections (as performed in other studies). Differences in mouse surgeon, histologic processing, and animal housing could also contribute to variability in experimental measurements. Despite this, the increase in LP thickness between injured and control airways with this model appears reproducible and comparable across institutions.

### Interpretation in the Context of the Totality of Evidence

These findings add to and extend an emerging body of literature demonstrating that, rather than serving solely as passive conduits for interstitial fluid drainage, lymphatic vessels actively regulate tissue repair throughout the body. In the lungs, impaired lymphatic drainage in inflammatory airway disease results in interstitial fluid overload, bronchial edema and airway obstruction^12,25,26^. As a case in point, pulmonary lymphatic drainage has been shown to be necessary for inflammation resolution in a murine model of Acute Respiratory Distress Syndrome, and treatment with VEGF-C156S (a VEGFR-3-specific synthetic VEGF-C analogue) was shown to be sufficient to rejuvenate lymphatics and promote resolution of inflammation in the short term in the lung^9^. The development of therapies that promote organized lymphangiogenesis in the airway remains an active area of investigation and a promising research frontier^27^.

Interestingly, lymphatic dysfunction is an under-appreciated feature of many established risk factors for LTS, including diabetes, smoking, and obesity in adults, as well as prematurity, low birthweight, and congenital heart disease in neonates and children^1,8,28–36^. In diabetes, the molecular mechanisms underpinning lymphatic dysfunction include impaired VEGFR3-mediated lymphangiogenesis, increased lymphatic permeability, and impaired lymphatic contractility^37–42^. Similarly, obesity causes negative effects through numerous mechanisms including impairing lymphatic valve function and pumping as well as through perilymphatic accumulation of inflammatory cells and lipids^42–44^. Preterm infants also have documented disordered lymphatic drainage, likely due to abnormal ex utero airway development, though the precise pathophysiology remains poorly defined^30^. Thus, while often clinically invisible, inadequate lymphatic function and the resultant stasis of interstitial inflammatory fluid may be a common contributing pathway contributing to persistent airway inflammation and pathogenic remodeling across these clinical populations.

Numerous aspects of critical illness and clinical care affecting both adults and children also negatively impact airway lymphatics. For example, positive pressure ventilation, often necessary in the intensive care unit for acute and chronic respiratory failure, impairs lymphatic drainage by inverting the physiologic respiratory cycle in which negative intrathoracic pressure provides a favorable gradient for thoracic duct filling during inspiration which is then emptied during expiration. Instead, positive intrathoracic pressure increases intrathoracic interstitial pressure and central venous pressure, acting to collapse lymphatic vessels and reduce the pressure gradient that ordinarily drives thoracic duct outflow^45–50^. Other clinical factors with known, often deleterious impacts on lymphatic function include tissue hypoxia and hypoperfusion, sepsis, patient immobility, surgery, aspiration, and hyperoxia^9,51–56^. Although these associations remain speculative in the context of airway injury, our data suggest that diminished lymphatic reserve may represent a common biological pathway through which otherwise disparate clinical conditions predispose patients to abnormal airway healing. Enhancing existing clinical care protocols with an eye toward supporting lymphatic function may have a role in improving airway outcomes.

### Controversies Raised by This Study

First, the cellular mechanisms by which VEGFR-3 inhibition impairs airway repair remain undefined. Whether defective healing reflects impaired fluid clearance, altered immune cell trafficking, reduced antigen transport, or direct lymphatic-stromal cell signaling interactions requires further investigation.

Second, in many chronic inflammatory diseases, expansion of lymphatic vessels has traditionally been interpreted as evidence of disease progression. Our data support the alternative hypothesis that lymphatic remodeling represents an adaptive repair program that promotes restoration of homeostasis following acute airway damage and enables wound healing.

Finally, the relationship between early lymphatic dysfunction and later fibrotic remodeling remains unexplored. Longer observation periods may clarify how persistent lymphatic alterations influence chronic remodeling.

### Future Research Directions

Several avenues for future investigation emerge from this work. Mechanistically, defining the cell populations and inflammatory mediators regulated by airway lymphatics will be critical for understanding how lymphangiogenesis coordinates tissue repair. Single-cell transcriptomics, spatial profiling, and lineage-specific genetic models targeting lymphatic endothelial cells will likely provide key insights into the cellular interactions that govern airway healing in time and space. Equally important will be determining whether therapeutic enhancement of lymphatic function can improve outcomes after airway injury. Administration of VEGFR-3 agonists, strategies that augment lymphatic endothelial signaling, or localized delivery of lymphangiogenic therapies at the time of airway injury may represent novel approaches for preventing pathologic airway remodeling. Functional studies using human laryngotracheal stenosis specimens will be essential to validate how well the lymphatic responses identified in this murine model are conserved in human disease. New clinical diagnostic tools and clinical informatics-based approaches could be developed to identify patients at risk for lymphatic dysfunction who may benefit from targeted treatments.

Ultimately, these studies support a conceptual shift: lymphatics are indispensable active regulators of airway inflammatory resolution and tissue regeneration. Defining how lymphatic biology integrates immune regulation, homeostasis, and wound repair may open new avenues to prevent and treat LTS after airway injury.

## Acknowledgments

This work was supported by an American Society of Pediatric Otolaryngology Research Career Development Award (J.S.P.) and a Triological Society Career Development Award (J.S.P.). We acknowledge the Translational Pathology Shared Resource supported by NCI/NIH Cancer Center Support Grant P30CA068485 and the Shared Instrumentation Grant S10 OD023475-01A1 for the Leica Bond RX (Immunohistochemistry).

## Additional information

### Ethical approval and informed consent statements

All animal procedures were approved by the Vanderbilt Institutional Animal Care and Use Committee and adhered strictly to institutional and U.S. national guidelines for animal welfare and research ethics.

### Data availability statement

The datasets generated during and/or analyzed during the current study are available from the corresponding author on reasonable request.

